# DeepEpitope: Leveraging Transformation-Based protein Embeddings for Accurate linear Cancer B-cell Epitope Identification

**DOI:** 10.1101/2025.05.23.655884

**Authors:** T Dhanushkumar, Pasupuleti Visweswara Rao, Karthick Vasudevan

## Abstract

Conventional cancer treatments tend to have serious side effects, leading to the quest for safer and more specific treatment modalities. Immunotherapy with vaccines has appeared as a promising option, with B-cell epitopes being crucial for the generation of humoral immunity. But identification of the correct B-cell epitopes of cancer is a severe challenge since current tools are not pre-trained with cancer-generated datasets. To bridge this gap, we introduce DeepEpitope, a command-line tool based on deep learning designed exclusively for the prediction of linear B-cell epitopes from cancer antigens. We compiled a high-quality dataset from the Cancer Epitope Database and used Evolutionary Scale Modeling (ESM) embeddings to represent epitope and non-epitope sequences as vectors of 1280 dimensions. These embeddings were employed to train five machine learning models (Logistic Regression, Random Forest, XGBoost, LightGBM, and Naive Bayes) and three deep learning models (Multilayer Perceptron [MLP], Convolutional Neural Network, and Bidirectional LSTM). Of these, the MLP model performed best with an AUC of 0.85 and a benchmark AUC of 94%. In comparison with other tools like BepiPred (60%) and LBtope (54%), DeepEpitope demonstrated much higher predictive accuracy. It is a Linux-based command-line tool that can be accessed for free at: https://github.com/karthick1087/DeepEpitope.

## Introduction

Cancer remains the leading cause of morbidity and mortality worldwide despite being a major area of research [1]. The complexity and heterogeneity of cancer have necessitated the development of diverse treatment options to manage and potentially cure the disease at different stages. Traditional cancer treatments include surgical removal of the tumor, followed by chemotherapy and radiotherapy. More recently, hormonal therapy targeting hormone-sensitive cancers, such as certain breast and prostate cancers, has been developed by blocking hormone production or interfering with hormone action [2]. However, all of these traditional therapies can be painful and have significant side effects. Therefore, researchers are exploring more effective strategies, such as immunotherapy. Cancer vaccines have emerged as a promising addition to cancer treatment options [3].

Cancer vaccines represent a promising approach for preventing and treating various types of cancer by harnessing the body’s immune system. These vaccines have demonstrated varying degrees of success in clinical trials and real-world applications [4]. Preventive vaccines have shown remarkable efficacy, such as an 89% reduction in cervical intraepithelial neoplasia (CIN) grade 3 when administered with the Human Papillomavirus (HPV) vaccine, which protects against HPV infections. This HPV vaccine has also shown positive outcomes in preventing head and neck cancers, anal cancer, penile cancer, vaginal cancer, and vulvar cancer by blocking the viral infections that can lead to these malignancies [5].

The FDA has approved therapeutic cancer vaccines, such as Sipuleucel-T (Provenge®) for prostate cancer. This personalized vaccine involves removing some of the patient’s immune cells, exposing them to prostate cancer antigens, and then reinfusing them back into the body. Studies have shown that Sipuleucel-T extends survival in patients with metastatic prostate cancer [6]. Epitope-based vaccines represent a sophisticated approach to cancer vaccination by targeting specific molecular structures (epitopes) recognized by the immune system. For example, the MAGE-A3 vaccine, which targets the MAGE-A3 antigen expressed in non-small cell lung cancer (NSCLC), showed a 27% improvement in phase 2 clinical trials [7]. GVAX-NSCLC, another therapeutic vaccine, demonstrated encouraging results in a Phase I/II study for NSCLC [8]. HER-Vaxx, a gastric cancer vaccine, achieved an 89.7% disease-free survival (DFS) rate compared to 80.2% in controls in phase 2 trials [9]. Additionally, mRNA-4157 (Moderna), a melanoma vaccine, reduced the risk of recurrence or death by 44% (hazard ratio = 0.56) in patients with resected stage III/IV melanoma [10]. Epitope-based vaccines incorporate T-cell and B-cell epitopes with adjuvants. T cell and B cell epitopes are the specific regions of tumor antigens that are recognized by the T and B cells. B-cell epitopes play a pivotal role in vaccines by eliciting a robust humoral immune response. B cells act as antigen-presenting cells (APCs), processing and presenting tumor antigens to helper T cells, thereby bridging innate and adaptive immunity [11]. The strategic inclusion of B-cell epitopes in cancer vaccines has shown promise in enhancing vaccine efficacy in many clinical trials, for instance, HER-Vaxx, which showed 89.7% DFS, used 3 B-cell epitopes in it [9].

One of the major challenges in developing epitope-based cancer vaccines is the accurate identification and incorporation of relevant epitopes [12]. To address this limitation, we designed DeepEpitope, a cancer-specific B-cell epitope prediction tool that focuses on predicting linear epitopes. Currently, no tools specifically tool that is trained with cancer epitopes exist, researchers often rely on tools such as ABCpred [13], BCPred [14], BepiPred-2.0 [15], and other tools for vaccine design and epitope prediction. These tools are trained on diverse epitope datasets from multiple diseases and typically utilize features like physicochemical properties and dipeptide sequences of epitopes [16]. Most existing tools are based on datasets from the Immune Epitope Database (IEDB), which contains a vast and heterogeneous collection of epitopes spanning numerous diseases [17]. More recently, advanced tools like BepiPred-3.0 [18] and CALIBER [19] have incorporated embedding techniques such as Evolutionary Scale Modeling (ESM), a protein language model developed by Meta AI (Facebook AI Research) that leverages transformer-based deep learning architectures, similar to NLP (Natural Language Processing) models like BERT (Bidirectional Encoder Representations from Transformers) and GPT (Generative Pre-trained Transformer), but trained on protein sequences [20].

In our work, we utilized the ESM-2 (Evolutionary Scale Mode). model, which encodes epitope sequences into a 1280-dimensional vector representation. Leveraging this rich vector information, we developed DeepEpitope using a curated dataset of epitopes sourced specifically from the cancer epitope database, enabling a more focused and accurate prediction of tumor-specific B-cell epitopes, a detailed workflow followed for this study is illustrated in Figure 1.

**Figure 1.**
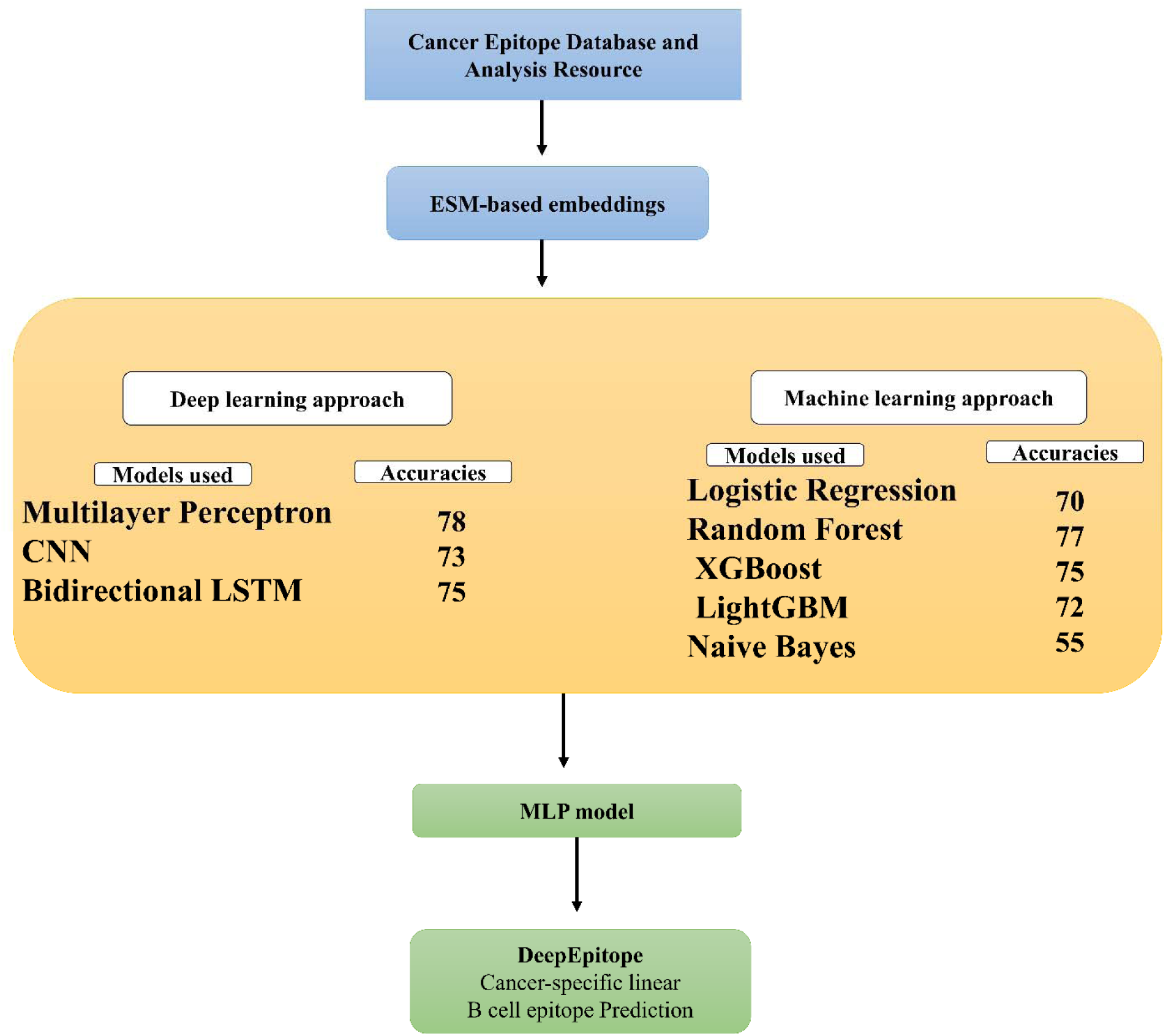
Overall workflow of this study

## Results

### Performance of Individual ML/DL Models

For this study, we implemented five machine learning (ML) and three deep learning (DL) models, including Logistic Regression, Random Forest, XGBoost, LightGBM, Naïve Bayes, Multilayer Perceptron (MLP), Convolutional Neural Network (CNN), and Bidirectional LSTM (BiLSTM). These models were trained and validated using a curated dataset from the Cancer Epitope Database and Analysis Resource (CEDAR) [21] focused on linear B-cell epitopes derived from tumor antigens, and training was done by transforming the epitope information into to 1280 D vector using the ESM2 embedding technique. To evaluate the predictive capabilities of the trained models, model’s performance was evaluated using the following analysis.

### ROC and PR Curve Analysis

Receiver Operating Characteristic (ROC) curve analysis (Figure 2) revealed that deep learning models outperformed traditional ML models. MLP achieved the highest AUC score of 0.85, followed closely by BiLSTM (AUC = 0.83) and CNN (AUC = 0.82), outperforming classical algorithms like Random Forest (0.78), Logistic Regression (AUC = 0.75), and Naïve Bayes (AUC = 0.63). This indicates a higher true positive rate at lower false positive rates for DL based approaches. Precision-Recall (PR) curves (Figure 3) further reinforced these results, with MLP attaining the highest precision of 0.80, while CNN and BiLSTM followed closely with a precision of 0.79. These high-precision values demonstrate the models’ ability to reduce false-positive predictions in imbalanced datasets, a crucial factor for reliable epitope prediction.

**Figure 2.**
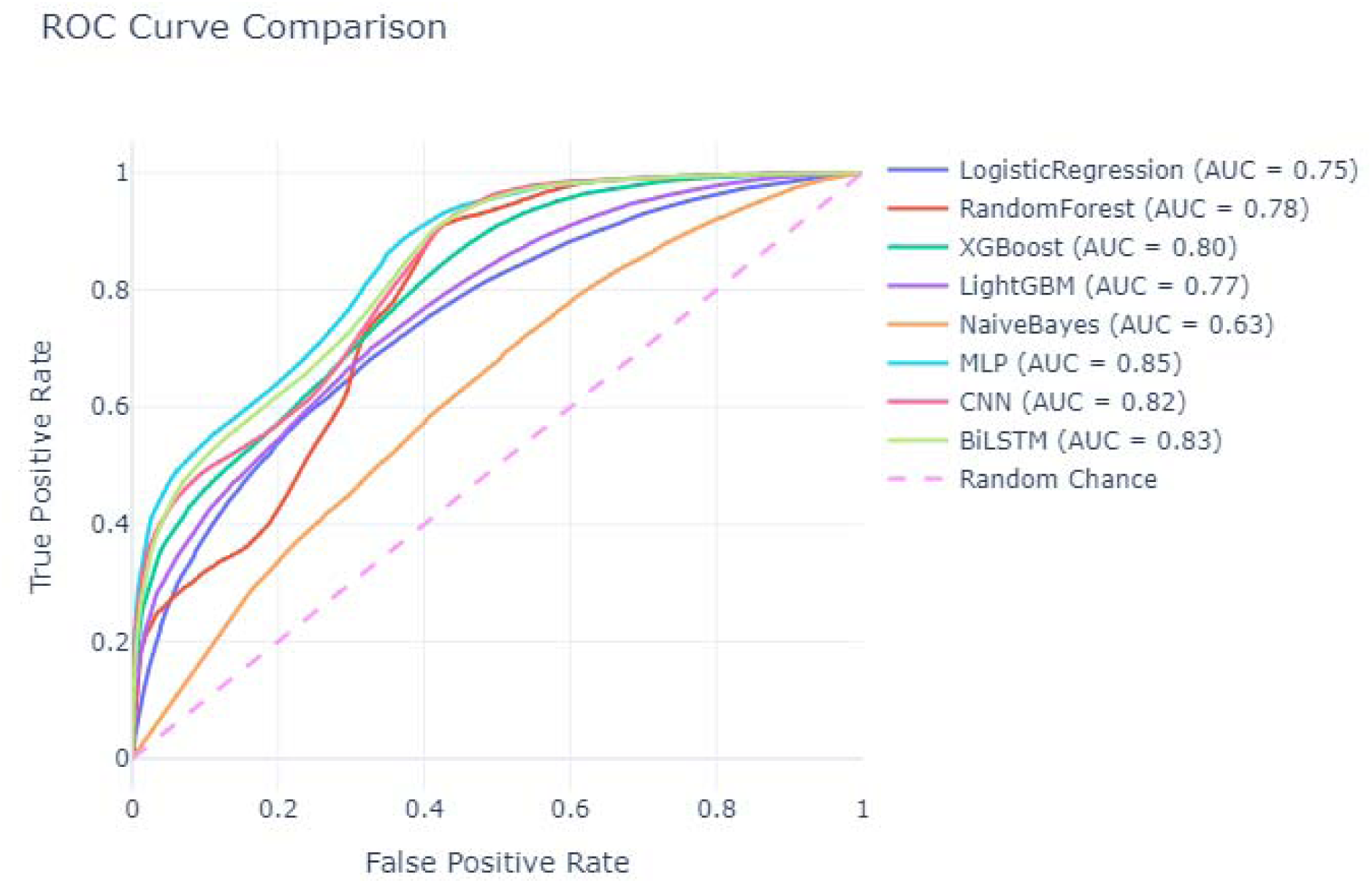
Receiver Operating Characteristic (ROC) curves for individual machine learning and deep learning models.

**Figure 3.**
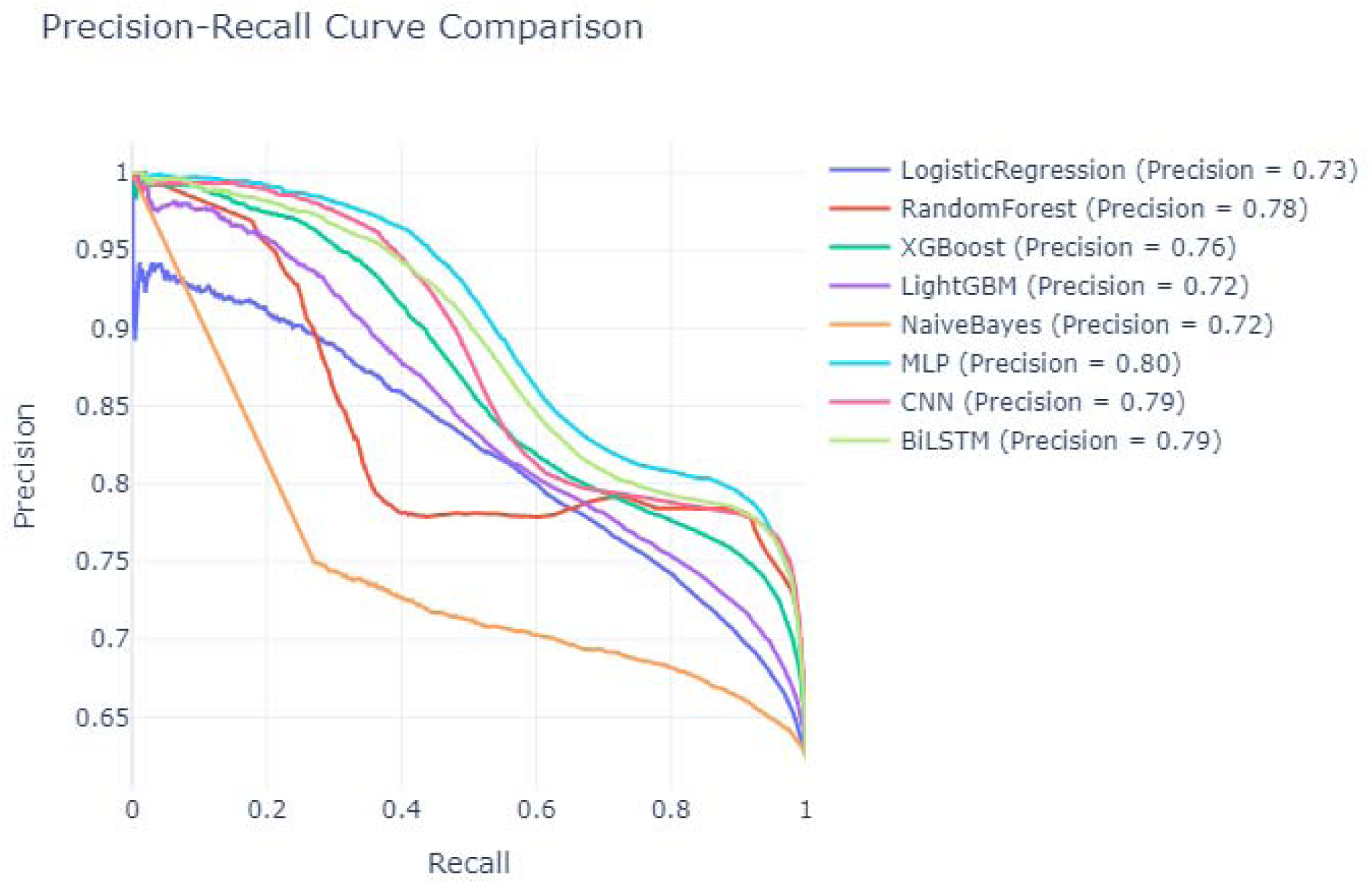
Precision-Recall (PR) curves of machine learning and deep learning models.

### Accuracy, Calibration Analysis, Parallel Coordinate Plot, Probability Distribution analysis

Model-wise accuracy comparison (Figure 4) revealed that MLP achieved the highest accuracy (78.0%), followed by Random Forest (76.6%) and BiLSTM (75.3%). Naïve Bayes exhibited the poorest performance with an accuracy of 54.7%, reflecting its limited ability to generalize over the dataset. Calibration curves (Figure 5) indicated that the predicted probabilities of the MLP, CNN, and BiLSTM models are better aligned with true outcome frequencies compared to traditional models. MLP and CNN predictions closely tracked the ideal calibration line, suggesting high reliability in probability estimates, a critical aspect in downstream tool development and output representation. The parallel coordinates plot (Figure 6) highlighted MLP performer across all key metrics Accuracy (0.7797), Precision (0.803), Recall (0.8975), F1-score (0.8287), and AUC (0.8478). Violin plots of predicted probability distributions (Figure 7) showed that MLP, BiLSTM, and CNN generated sharper and well-distributed probability estimates, supporting the robustness of their confidence in class assignment. In contrast, Naïve Bayes showed a flatter and less distinctive distribution, reinforcing its lower performance.

**Figure 4.**
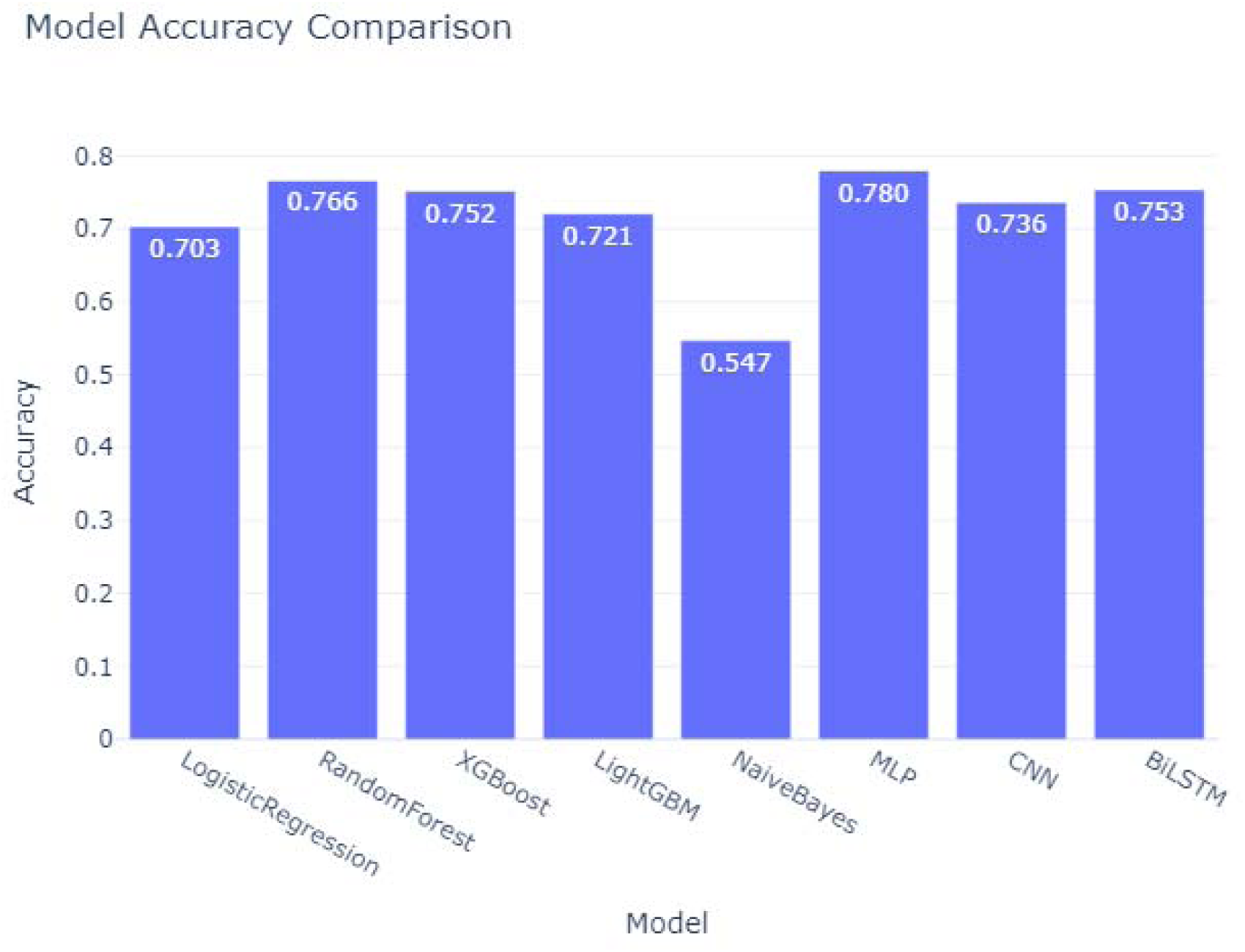
Accuracy comparison of machine learning and deep learning models.

**Figure 5.**
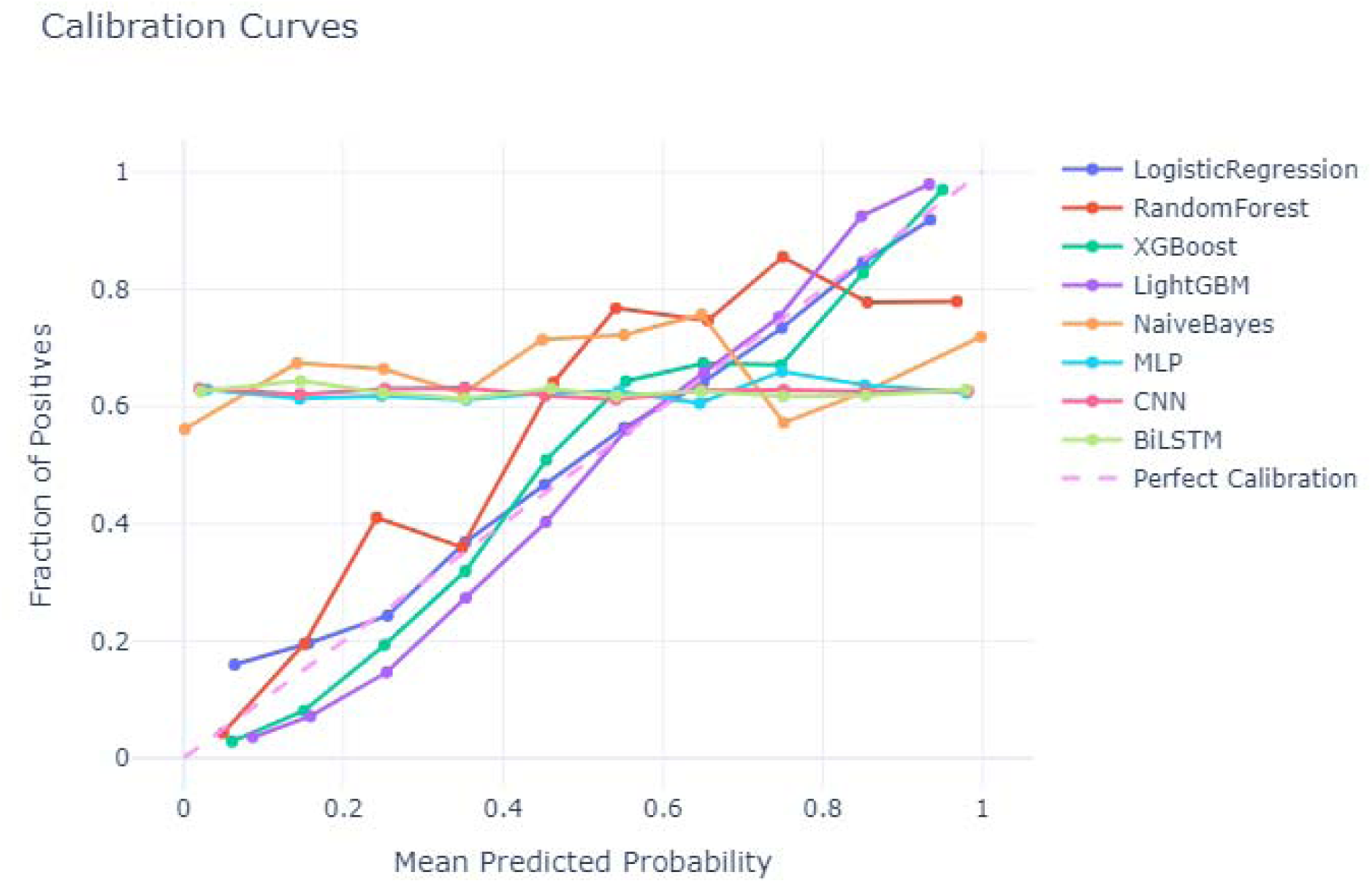
Calibration curves for all models

**Figure 6.**
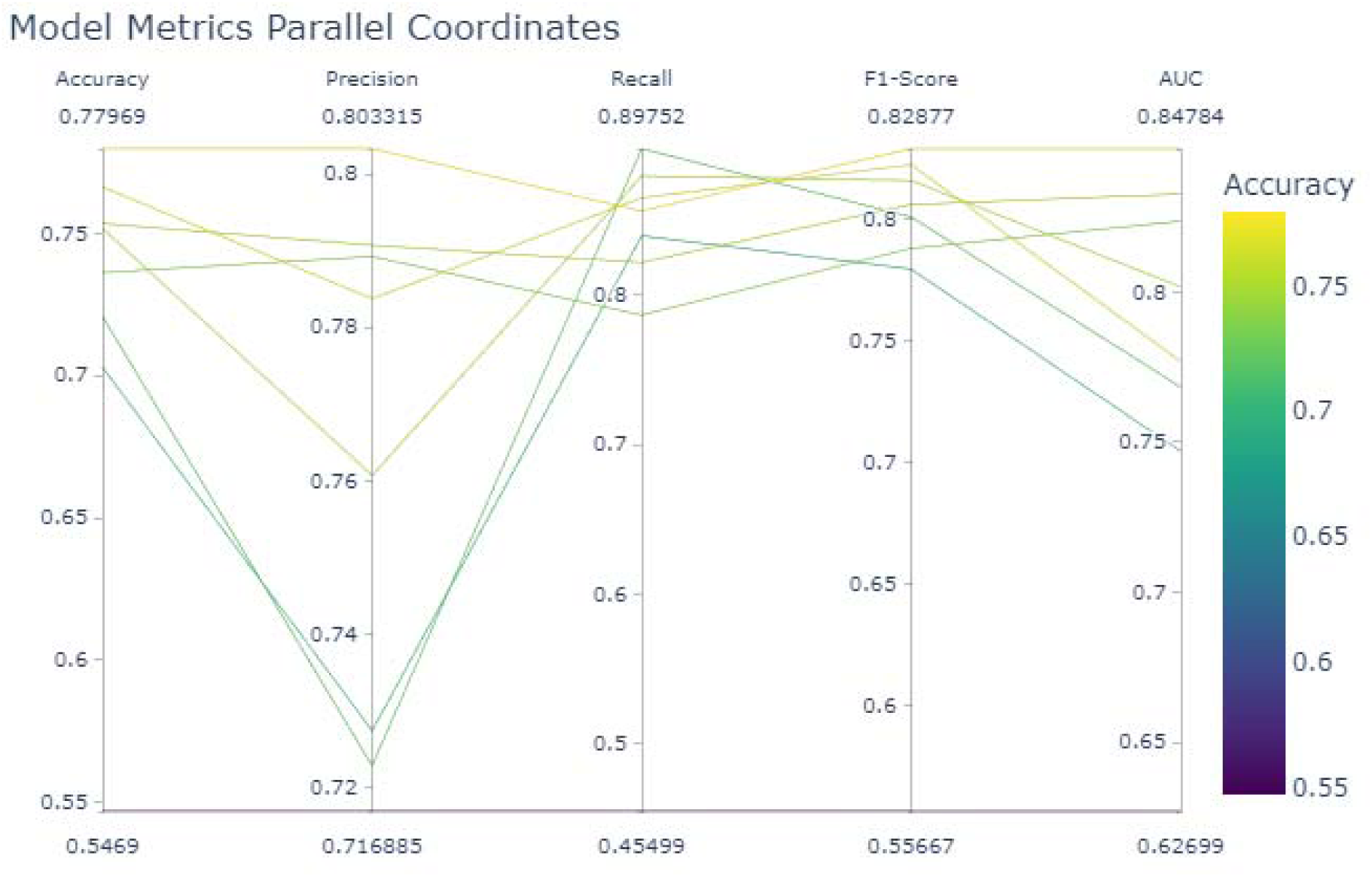
A parallel coordinates plot summarizing model performance across multiple metrics for the MLP model.

**Figure 7.**
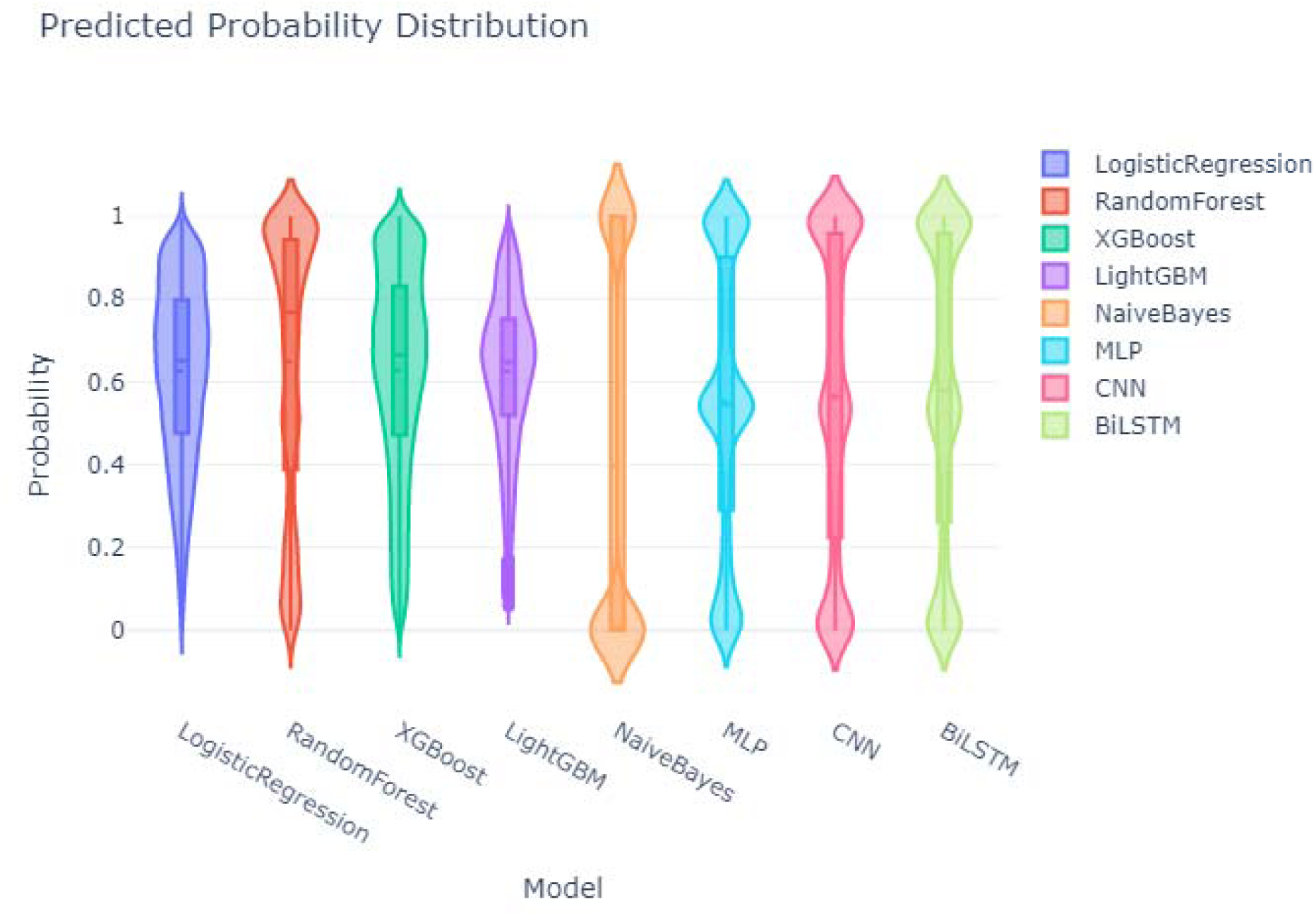
Violin plots of predicted probability distributions.

### Command-Line-based Tool deployment

Overall evaluation demonstrated that deep learning-based models, particularly MLP showed superior performance over traditional ML methods, achieving the highest predictive accuracy. We selected the MLP model for command-line tool development. To ensure broad usability, we implemented the tool as a Linux-based command-line utility, hosted on GitHub, allowing seamless local execution and integration into high-throughput bioinformatics pipelines.

Unlike many existing web-based epitope prediction tools that process only single sequences or lack flexibility in input customization, DeepEpitope accepts multi-FASTA files and provides user defined control over peptide length by mentioned minimum and maximum lengths and epitope selection (e.g., threshold, top rankings). Additionally, many publicly available tools are unable to handle full-length proteins. Our approach addresses these limitations by supporting full-length input proteins and customizable output formats, offering a more robust and scalable solution for cancer specific B cell epitope predations.

### Comparison of the designed model with existing tools and validation

To independently validate the performance of DeepEpitope, we curated a benchmark dataset consisting of 45 cancer-related B-cell epitopes and 31 non-epitopes from the Immune Epitope Database (IEDB). All selected peptides were strictly excluded from the model’s training set to ensure unbiased evaluation. The predictive performance of DeepEpitope was then compared against two widely used linear B-cell epitope prediction tools: BepiPred-2.0 [16] and LBtope [22]. Figures 8A and 8B depict the Precision-Recall (PR) and Receiver Operating Characteristic (ROC) curves, respectively, highlighting the comparative performance of the models. DeepEpitope outperformed both traditional tools across all metrics

**Figure 8.**
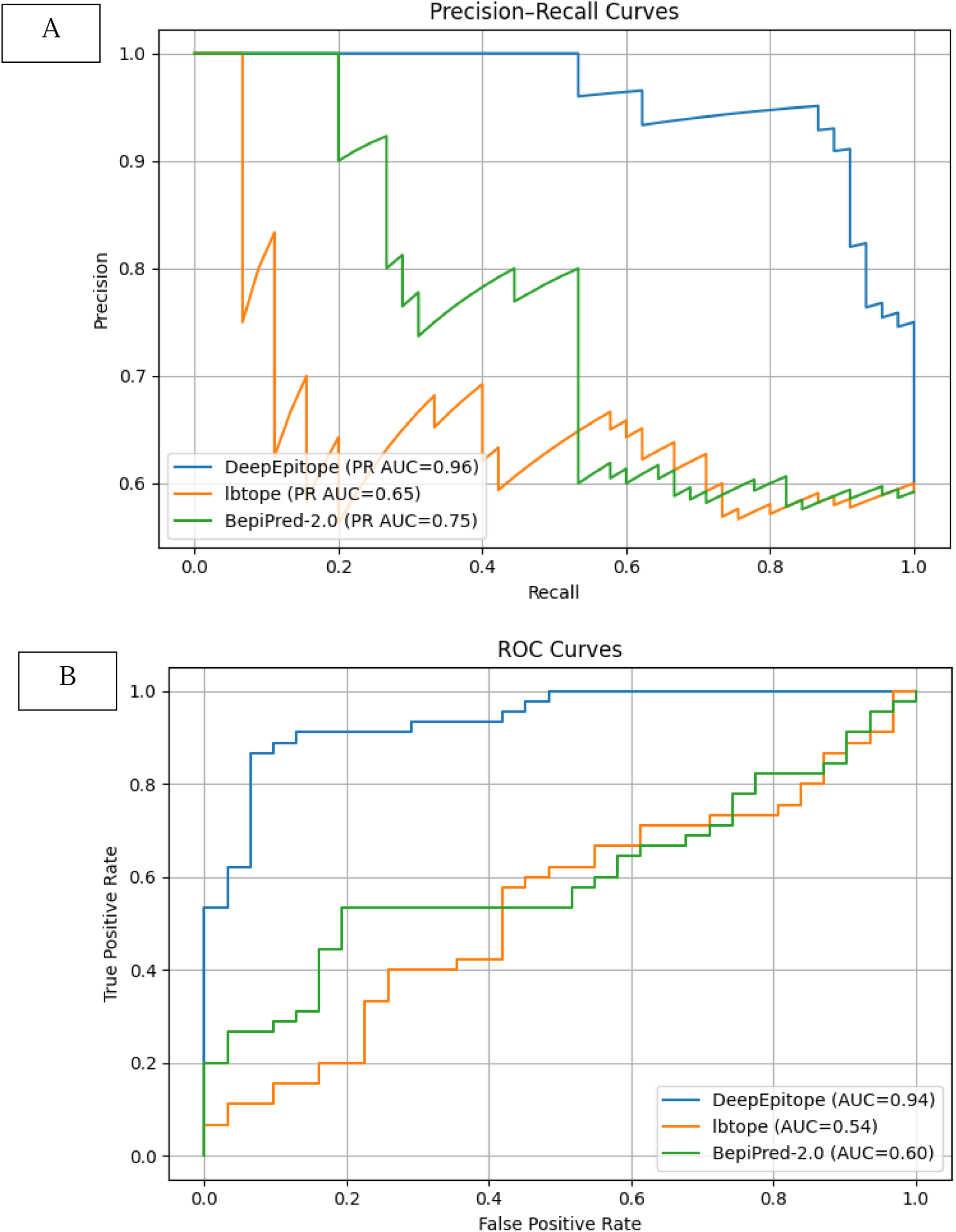
A) Precision-Recall (PR) curve comparing DeepEpitope with existing tools, B) ROC curve comparing DeepEpitope with existing tools.

In the PR curve (Figure 8A), DeepEpitope achieved a PR AUC of 0.96, significantly higher than BepiPred-2.0 (0.75) and LBtope (0.65). This indicates superior performance in distinguishing epitopes from non-epitopes, particularly in datasets with class imbalance. In the ROC analysis (Figure 8B), DeepEpitope reached an AUC of 0.94, while BepiPred-2.0 and LBtope achieved 0.60 and 0.54, respectively. This demonstrates DeepEpitope’s stronger ability to maximize true positive rates while minimizing false positives. These results validate that DeepEpitope generalizes well to unseen cancer-related epitope sequences and outperforms existing state-of-the-art tools. The substantial margin in both AUC-ROC and AUC-PR underscores the value of leveraging transformer-based embeddings and deep learning architectures for robust linear epitope prediction. Benchmark data used in this analysis and each tool’s scores and output can be found in the supplementary file.

## Discussion

In this study, we designed DeepEpitope, a command-line-based computational tool dedicated to the prediction of linear B-cell epitopes, specifically from cancer-related antigens. The main objective of this study was to address the current limitations in B-cell epitope prediction tools and to deliver a model capable of accurately identifying epitopes derived from tumor-associated antigens using advanced deep learning and feature embedding techniques [23].

B-cell epitope prediction remains one of the most crucial yet challenging components of immunoinformatics. B-cell epitopes are antigenic regions recognized by antibodies, and accurate identification of such epitopes plays a pivotal role in vaccine design, antibody engineering, and immunotherapy development [24]. However, existing epitope prediction tools face significant limitations. Many tools, such as BepiPred-2.0 [16] and LBtope [22], were trained on broad or general datasets not specifically curated for cancer-related antigens. As a result, their predictive performance is not up to the mark when applied to tumor antigens, which possess distinct immunological and biochemical properties compared to microbial or viral antigens. Another major drawback of current tools is that none are trained to predict epitopes specific to cancer, nor do they leverage cancer epitope datasets during training, which creates a major gap [25].

B-cell epitopes are broadly categorized into linear (continuous) and conformational (discontinuous) epitopes. in this study, we are focusing on linear B-cell epitopes, which are continuous sequences of amino acids on a tumor antigen of cancer [26].

Feature engineering is the core of the machine learning (ML) or deep learning (DL) pipeline in biological sequence analysis. Traditional approaches to epitope prediction have relied upon features like amino acid composition, physicochemical properties, or secondary structure elements, atomic count like nitrogen, hydrogen, oxygen, and carbon counts, and dipeptide composition (400 features) [27]. However, these features cannot often capture the contextual, hierarchical, and evolutionary information embedded in protein sequences. And offer very limited features to train the models, which may result in wrong predictions

To address this, we employed ESM2, a transformer-based protein language model developed by Meta AI. ESM2 is trained on massive protein sequence datasets and can embed protein sequences into high-dimensional vectors that retain meaningful biochemical and structural context. By converting each peptide into a 1280-dimensional embedding, we ensured that our models received a rich and highly accurate representation of cancer epitopes. This choice plays a very important role in achieving high accuracy in our models [28].

Using these high-quality embeddings, we trained a suite of eight supervised learning models, consisting of five ML algorithms. Logistic Regression, Random Forest, Naïve Bayes, XGBoost, and LightGBM, and three DL models, which include Multilayer Perceptron (MLP), Convolutional Neural Network (CNN), and Bidirectional LSTM (BiLSTM).

Among ML models, Random Forest and XGBoost showed relatively strong performance with accuracy rates of 0.76 and 0.75, however, Logistic Regression, LightGBM, and Naïve Bayes showed optimal performance. DL models, particularly MLP and BiLSTM, showed superior performance. Through a detailed comparative analysis, MLP emerged as the top-performing model, achieving the highest scores across multiple metrics, including ROC-AUC (0.85), PR-AUC (0.80), Accuracy (78.0%), and F1-score (0.83). Precision-recall analysis further emphasized MLP’s ability to maintain high precision while minimizing false positives, crucial in datasets with imbalanced class distributions. Calibration plots and violin plots of predicted probabilities confirmed MLP’s robustness and reliability in confidence estimation. BiLSTM and CNN followed closely in terms of performance, demonstrating that deep learning architectures offer distinct advantages over traditional ML models for complex bioinformatics tasks such as epitope prediction. However, MLP’s simplicity, training stability, and superior overall accuracy made it the ideal candidate for deployment in a command-line tool [29].

To validate DeepEpitope’s practical performance and real-world applicability, we benchmarked it against two widely used tools, BepiPred-2.0 and LBtope, using an external test set consisting of experimentally validated cancer epitopes and non-epitopes curated from the Immune Epitope Database (IEDB). These benchmarks revealed that DeepEpitope significantly outperforms existing tools, achieving an ROC-AUC of 0.94 and a PR-AUC of 0.96, compared to much lower values obtained by the competing tools. However, these two tools were solely built using IEDB datasets, which contain a diverse set of epitopes, which may be a reason for this low performance. Through this analysis, it is clear that our tool predicts the cancer epitope and non-epitope. These findings underscore the importance of using high-quality embeddings (like ESM2) and cancer-specific data in model training.

Importantly, DeepEpitope is a fully functional command-line tool designed for high-throughput, reproducible, and scalable analysis. Users can run the tool with a single command, supplying a multi-FASTA file or a single FASTA sequence in FASTA form of antigen sequences, and receive as output a detailed list of predicted linear B-cell epitopes (all output predictions and the top 20 epitopes). The tool is engineered to support additional options, such as adjusting peptide length windows, predicting full-length peptides to classify whether the given input is an epitope or non-epitope, setting prediction thresholds, choosing top-n peptides based on scores, and exporting results in user-friendly formats (CSV or Excel). These capabilities make DeepEpitope highly adaptable for integration into automated bioinformatics pipelines.

DeepEpitope also demonstrates the power of transfer learning and pre-trained protein language models in downstream tasks. ESM2 embeddings eliminate the need for traditional features, enabling models to train with semantically rich input vectors that capture sequence-level and evolutionary context. This approach significantly lessens the burden of manual feature extraction while allowing the application of advanced learning techniques.

Despite these advances, there are some limitations to consider. The model currently focuses solely on linear epitopes and does not address conformational epitopes, which constitute a significant portion of antibody-targeted regions. In the future, integrating structural data or using 3D-aware embeddings could enhance the tool’s scope. Additionally, experimental validation has to be done with the prediction from this tool.

## Conclusion

DeepEpitope provides a powerful and precise deep learning-based predictor of linear B-cell epitopes of cancer antigens. The integration of cancer-specific training data with ESM-2 protein embeddings makes it perform notably better than available tools in terms of prediction capability. The use of MLP-based architecture facilitates fast, high-throughput analysis via a user-controllable command-line interface, rendering it amenable for incorporation into vaccine design pipelines. Although presently restricted to linear epitopes, DeepEpitope fills an important void in cancer immunoinformatics and paves the way for future expansion to include conformational epitope prediction as well as experimental validation.

## Methods and Methodology

### Data Collection and Preprocessing

In this study, we utilized the CEDAR as the primary source for our dataset. CEDAR is a comprehensive, freely accessible repository that curates cancer-specific epitope and receptor data from the literature. It provides detailed information on various aspects of cancer epitopes, including their molecular characteristics, associated metadata, and experimental validation methods. The raw data files from CEDAR encompass extensive details such as the authorship, publication titles, experimental assays performed, and the classification of epitopes as either linear or conformational [21]. Additionally, the dataset includes information on the parent tumor antigens from which these epitopes are derived, along with other relevant immunological assays and other contexts. For our analysis, we focused exclusively on linear B-cell epitopes. From the raw dataset, we extracted entries labeled as positive (epitopes) and negative (non-epitopes). To ensure data quality and relevance, we performed several preprocessing steps like Deduplication (removing duplicate entries to prevent redundancy), Filtering Non-natural Epitopes (Excluded epitopes containing non-natural amino acids or modifications that could introduce bias or are not representative of typical biological sequences) After these preprocessing steps, we obtained a balanced dataset comprising 37,609 linear B-cell epitopes and 40,637 non-epitopes. This substantial and balanced dataset

### Feature Extraction Using ESM2

To effectively capture the complex biochemical and structural characteristics of cancer-related linear B-cell epitopes, we employed ESM2, a state-of-the-art transformer-based protein language model developed by Meta AI’s Fundamental AI Research team. ESM2 is trained on extensive protein sequence datasets and is designed to generate high-dimensional embeddings that encapsulate rich contextual information about protein sequences [20].

### Model Development

To build a robust and accurate prediction system for cancer-specific linear B-cell epitopes, we adopted a dual-strategy approach involving both traditional Machine Learning (ML) algorithms and modern Deep Learning (DL) architectures. This hybrid strategy was aimed at leveraging the interpretability of ML models and the expressive learning capabilities of DL models to assess their relative performance and choose the most effective one for deployment.

We implemented five supervised ML classifiers. Logistic Regression (LR), a linear model used for binary classification. It was configured with max iterations of 5000 and solver=’lbfgs’ to ensure convergence on high-dimensional ESM2 embeddings. Random Forest (RF) is an ensemble-based decision tree classifier using n_estimators=100, suitable for non-linear data and offering feature importance analysis. XGBoost, a gradient boosting framework known for its superior speed and performance, is initialized with eval_metric=’logloss’ to optimize binary classification tasks. LightGBM (LGBM) is another gradient boosting method optimized for performance on large datasets. It provides fast training speed and low memory usage. Naïve Bayes (NB) is a probabilistic classifier based on Bayes’ theorem. These models were trained on the 1280-dimensional ESM2 feature embeddings. A standard 80-20 stratified train-test split was used to preserve the class balance in both training and testing sets. Each model was fitted using the training set and evaluated on the held-out test set [30].

To fully exploit the representational power of ESM2 embeddings, we also implemented three DL models using PyTorch. A Multilayer Perceptron (MLP) was trained on the data with three fully connected layers (1280 → 512 → 128 → 1) with batch normalization and dropout (0.4) after each layer to prevent overfitting. MLP served as the baseline deep learning model due to its simplicity and strong generalization ability in feedforward networks. Similarly, a Convolutional Neural Network (CNN) was trained with two 1D convolutional layers (Conv1D → MaxPooling → Conv1D) followed by two fully connected layers. Batch normalization and dropout were used to regularize learning. CNNs are effective in capturing local feature patterns and interactions between adjacent amino acid embeddings, useful for sequence classification tasks. Finally, Bidirectional Long Short-Term Memory (BiLSTM) was trained with a bidirectional LSTM layer (hidden_dim = 64) followed by a fully connected layer. Each sample was passed as a sequence of one timestep to match the LSTM format. BiLSTM was used to capture long-term dependencies and directional context, especially useful in biological sequences where motif positioning can influence function [31–32].

### Performance Evaluation Metrics

For the model evaluation, we employed a comprehensive set of evaluation metrics to assess classification performance. These included accuracy, precision, recall, F1-score, ROC-AUC, PR-AUC, and comparative matrices. Accuracy provided a general measure of correctness, while precision and recall offered deeper insight into the balance between false positives and false negatives, which is especially important in biological datasets. The F1-score, being the harmonic mean of precision and recall, was particularly valuable in evaluating model robustness when classes were closely matched. ROC and precision-recall curves further helped visualize how well models separated the two classes across different thresholds. In addition, a classification report provided a detailed summary of performance across classes. Calibration curves were also generated to assess how well the predicted probabilities reflected actual outcomes, an important aspect for clinical or therapeutic decision-making. Finally, a parallel coordinates plot summarizes model performance across multiple metrics [34].

## Data availability

The tool is publicly available at https://github.com/karthick1087/DeepEpitope.

## CRediT authorship contribution statement

Dhanushkumar T: Writing – review & editing, Writing – original draft, Codes writing and editing, Visualization, Formal analysis, Data curation. Pasupuleti Visweswara Rao: Writing – review & editing, Writing – original draft. Karthick Vasudevan: Writing – review & editing, Writing – original draft, codes writing and editing, Visualization, Formal analysis, Conceptualization, project administration, Supervision.

## References

1. Pucci, C., Martinelli, C., & Ciofani, G. (2019). Innovative approaches for cancer treatment: Current perspectives and new challenges. ecancermedicalscience, 13, 961.

2. Debela, D. T., Muzazu, S. G., Heraro, K. D., Ndalama, M. T., Mesele, B. W., Haile, D. C., … & Manyazewal, T. (2021). New approaches and procedures for cancer treatment: Current perspectives. SAGE open medicine, 9, 20503121211034366.

3. Zugazagoitia, J., Guedes, C., Ponce, S., Ferrer, I., Molina-Pinelo, S., & Paz-Ares, L. (2016). Current challenges in cancer treatment. Clinical therapeutics, 38(7), 1551–1566.

4. Saxena, M., van der Burg, S. H., Melief, C. J., & Bhardwaj, N. (2021). Therapeutic cancer vaccines. Nature Reviews Cancer, 21(6), 360–378.

5. Cai, X., & Xu, L. (2024). Human Papillomavirus-Related Cancer Vaccine Strategies. Vaccines, 12(11), 1291. 10.3390/vaccines12111291

6. Cheever, M. A., & Higano, C. S. (2011). PROVENGE (Sipuleucel-T) in prostate cancer: the first FDA-approved therapeutic cancer vaccine. Clinical Cancer Research, 17(11), 3520–3526.

7. McQuade, J. L., Homsi, J., Torres-Cabala, C. A., Bassett, R., Popuri, R. M., James, M. L., … & Hwu, W. J. (2018). A phase II trial of recombinant MAGE-A3 protein with immunostimulant AS15 in combination with high-dose Interleukin-2 (HDIL2) induction therapy in metastatic melanoma. BMC cancer, 18, 1–9.

8. Wang, X., Niu, Y., & Bian, F. (2025). The progress of tumor vaccines clinical trials in non-small cell lung cancer. Clinical and Translational Oncology, 27(3), 1062–1074.

9. Tobias, J., Maglakelidze, M., Andrić, Z., Ryspayeva, D., Bulat, I., Nikolić, I., … & Wiedermann, U. (2024). Phase II Trial of HER-Vaxx, a B-cell Peptide-Based Vaccine, in HER2-Overexpressing Advanced Gastric Cancer Patients Under Platinum-Based Chemotherapy (HERIZON). Clinical Cancer Research, 30(18), 4044–4054.

10. Rojas, L. A., Sethna, Z., Soares, K. C., Olcese, C., Pang, N., Patterson, E., … & Balachandran, V. P. (2023). Personalized RNA neoantigen vaccines stimulate T cells in pancreatic cancer. Nature, 618(7963), 144–150.

11. Kaumaya, P. T. (2020). B-cell epitope peptide cancer vaccines: a new paradigm for combination immunotherapies with novel checkpoint peptide vaccine. Future Oncology, 16(23), 1767–1791.

12. Caoili, S. E. C. (2010). Benchmarking B cell epitope prediction for the design of peptide based vaccines: problems and prospects. BioMed Research International, 2010(1), 910524.

13. Malik, A. A., Ojha, S. C., Schaduangrat, N., & Nantasenamat, C. (2022). ABCpred: a webserver for the discovery of acetyl-and butyryl-cholinesterase inhibitors. Molecular Diversity, 1-21.

14. EL Manzalawy, Y., Dobbs, D., & Honavar, V. (2008). Predicting linear B cell epitopes using string kernels. Journal of Molecular Recognition: An Interdisciplinary Journal, 21(4), 243–255.

15. Jespersen, M. C., Peters, B., Nielsen, M., & Marcatili, P. (2017). BepiPred-2.0: improving sequence-based B-cell epitope prediction using conformational epitopes. Nucleic acids research, 45(W1), W24–W29.

16. Srivastava, K., & Srivastava, V. (2023). Prediction of conformational and linear B-cell epitopes on envelop protein of zika virus using immunoinformatics approach. International Journal of Peptide Research and Therapeutics, 29(1), 17.

17. Vita, R., Overton, J. A., Greenbaum, J. A., Ponomarenko, J., Clark, J. D., Cantrell, J. R., … & Peters, B. (2015). The immune epitope database (IEDB) 3.0. Nucleic acids research, 43(D1), D405–D412.

18. Clifford, J. N., Høie, M. H., Deleuran, S., Peters, B., Nielsen, M., & Marcatili, P. (2022). BepiPred 3.0: Improved B cell epitope prediction using protein language models. Protein Science, 31(12), e4497.

19. Israeli, S., & Louzoun, Y. (2024). Single-residue linear and conformational B cell epitopes prediction using random and ESM-2 based projections. Briefings in Bioinformatics, 25(2), bbae084.

20. Zhang, F., Li, J., Wen, Z., & Fang, C. (2024). FusPB-ESM2: Fusion model of ProtBERT and ESM-2 for cell-penetrating peptide prediction. Computational Biology and Chemistry, 111, 108098.

21. Koşaloğlu-Yalçın, Z., Blazeska, N., Vita, R., Carter, H., Nielsen, M., Schoenberger, S., … & Peters, B. (2023). The cancer epitope database and analysis resource (CEDAR). Nucleic acids research, 51(D1), D845–D852.

22. Singh, H., Ansari, H. R., & Raghava, G. P. (2013). Improved method for linear B-cell epitope prediction using antigen’s primary sequence. PloS one, 8(5), e62216.

23. El-Manzalawy, Y., & Honavar, V. (2010). Recent advances in B-cell epitope prediction methods. Immunome research, 6, 1–9.

24. Li, W. H., Su, J. Y., & Li, Y. M. (2022). Rational design of T-cell-and B-cell-based therapeutic cancer vaccines. Accounts of Chemical Research, 55(18), 2660–2671.

25. Sanchez-Trincado, J. L., Gomez-Perosanz, M., & Reche, P. A. (2017). Fundamentals and methods for T and B cell epitope prediction. Journal of immunology research, 2017(1), 2680160.

26. Van Regenmortel, M. H. (2009). What is a B-cell epitope?. Epitope Mapping Protocols: Second Edition, 3-20.

27. Liu, T., Shi, K., & Li, W. (2020). Deep learning methods improve linear B-cell epitope prediction. BioData mining, 13, 1–13.

28. Fan, W., Zhou, Y., Wang, S., Yan, Y., Liu, H., Zhao, Q., … & Li, Q. (2025). Computational Protein Science in the Era of Large Language Models (LLMs). arXiv preprint arXiv:2501.10282.

29. Nguyen, T. T., Nguyen, V. N., Tran, T. X., & Le, N. Q. K. (2024, November). Improved Linear B-Cell Epitope Prediction Using CNN and BiLSTM. In International Conference on Advances in Information and Communication Technology (pp. 466-475). Cham: Springer Nature Switzerland.

30. Kanoff, F. N. R. MACHINE LEARNING BASED TO PREDICT BCell EPITOPE REGION UTILIZING PROTEIN FEATURES.

31. Qi, Y., Zheng, P., & Huang, G. (2023). DeepLBCEPred: A Bi-LSTM and multi-scale CNN-based deep learning method for predicting linear B-cell epitopes. Frontiers in Microbiology, 14, 1117027.

32. Alghamdi, W., Attique, M., Alzahrani, E., Ullah, M. Z., & Khan, Y. D. (2022). LBCEPred: a machine learning model to predict linear B-cell epitopes. Briefings in bioinformatics, 23(3), bbac035.

33. Greenbaum, J. A., Andersen, P. H., Blythe, M., Bui, H. H., Cachau, R. E., Crowe, J., … & Peters, B. (2007). Towards a consensus on datasets and evaluation metrics for developing B cell epitope prediction tools. Journal of Molecular Recognition: An Interdisciplinary Journal, 20(2), 75–82.

34. Galanis, K. A., Nastou, K. C., Papandreou, N. C., Petichakis, G. N., & Iconomidou, V. A. (2019). Linear B-cell epitope prediction: a performance review of currently available methods. BioRxiv, 833418.

