## Supplementary Tables for "DeepEpitope: Leveraging Transformation-Based protein Embeddings for Accurate linear Cancer B-cell Epitope Identification"

Supplementary table 1. Benchmark data and their comparative probability scores in each tool with epitope information

| **Source Tumor antigen** | **IEDB link for epitope information** | **Peptide** | **True Label** | **DeepEpitope probability score** | **BepiPred-2.0 probability score** | **Lbtope probability score** |
| --- | --- | --- | --- | --- | --- | --- |
| Methionine aminopeptidase 1 | https://www.iedb.org/epitope/2258932 | AGISRELVDKLAAALE | 1 | 0.058966 | 0.4661 | 0.3297 |
| Solute carrier family 45 member 3 | https://www.iedb.org/epitope/1391841 | GASACDVSVRVVVGEP | 1 | 0.548672 | 0.466 | 0.3208 |
| TATA box-binding protein-associated factor RNA polymerase I subunit B | https://www.iedb.org/epitope/1311366 | GLKKKTILKKAGIGMCVKVSSIF  FINKQKP | 1 | 0.531142 | 0.527 | 0.6856 |
| Mucin-1 | https://www.iedb.org/epitope/1972710 | GVTSAPDTRPAPGSTAPPAH | 1 | 0.981875 | 0.3944 | 0.717 |
| Mucin-1 | https://www.iedb.org/epitope/2233080 | GVTSAPDTRPAPGSTAPPAHGVT | 1 | 0.996698 | 0.4098 | 0.6346 |
| ALK tyrosine kinase receptor | https://www.iedb.org/epitope/2257706 | GYQQQGLPLEAATAPGAGHYE  DTILKSKNSMNQPGP | 1 | 0.664547 | 0.4773 | 0.8632 |
| ALK tyrosine kinase receptor | https://www.iedb.org/epitope/2257707 | HKVHGSRNKPTSLWNPTYGSW  FTEKPTKKNNPIAKK | 1 | 0.982846 | 0.4815 | 0.5866 |
| Folate receptor alpha | https://www.iedb.org/epitope/2238360 | KDVSYLYRFNWNHCGEMA | 1 | 0.450629 | 0.437 | 0.5779 |
| Alpha-2-macroglobulin | https://www.iedb.org/epitope/2145013 | KMVSGFIPLKPTVKMLERSNH | 1 | 0.5655 | 0.442 | 0.5751 |
| Wilms tumor protein | https://www.iedb.org/epitope/2188420 | KRYFKLSHLQMHSRKH | 1 | 0.624728 | 0.5136 | 0.675 |
| Melanoma-associated antigen 3 | https://www.iedb.org/epitope/2275791 | KVAELVHFLLLKYRAREPVT | 1 | 0.10244 | 0.4545 | 0.3012 |
| Folate receptor alpha | https://www.iedb.org/epitope/2238388 | LGPWIQQVDQSWRKERV | 1 | 0.551308 | 0.4767 | 0.6768 |
| tyrosine-protein kinase erbB-2 | https://www.iedb.org/epitope/2237578 | LHCPALVTYNTDTFESMPN  PEGRYTFGASCV | 1 | 0.926623 | 0.4681 | 0.4949 |
| Endoplasmic reticulum chaperone BiP | https://www.iedb.org/epitope/2145052 | LIGRTWNDPSVQQDIKFL | 1 | 0.802472 | 0.4957 | 0.5496 |
| ALK tyrosine kinase receptor | https://www.iedb.org/epitope/2257714 | LLHVARDIACGCQYLEENHFI  HRDIAARNCLLTCPG | 1 | 0.822633 | 0.5047 | 0.5697 |
| Dickkopf-related protein 1 | https://www.iedb.org/epitope/2253758 | LNSNAIKNLPPPLGGAAG | 1 | 0.556962 | 0.4181 | 0.4649 |
| Heat shock factor protein 1 | https://www.iedb.org/epitope/2218374 | LSPQEPPRPIEAENSNPDSGKQ | 1 | 0.973673 | 0.4284 | 0.5869 |
| Heat shock factor protein 1 | https://www.iedb.org/epitope/2218376 | LSPQEPPRPPEAENSSPDSGKQ | 1 | 0.984235 | 0.4245 | 0.513 |
| ALK tyrosine kinase receptor | https://www.iedb.org/epitope/2257719 | LTANMKEVPLFRLRHFPCGNV  NYGYQQQGLPLEAAT | 1 | 0.18809 | 0.4699 | 0.4953 |
| Dickkopf-related protein 1 | https://www.iedb.org/epitope/2253763 | LYPGGNKYQTIDNYQPYP | 1 | 0.52698 | 0.4208 | 0.5113 |
| Heat shock factor protein 1 | https://www.iedb.org/epitope/2218381 | MDLAVGPGAAGPSNVPA | 1 | 0.934366 | 0.3982 | 0.4489 |
| Heat shock factor protein 1 | https://www.iedb.org/epitope/2218383 | MDLPVGPGAAGPSNVPA | 1 | 0.954167 | 0.3953 | 0.4003 |
| Syncytin-1 | https://www.iedb.org/epitope/243823 | NPSCPGGLGVTVCWTY | 1 | 0.701437 | 0.4106 | 0.629 |
| ALK tyrosine kinase receptor | https://www.iedb.org/epitope/2257722 | NTALPIEYGPLVEEEEKVPV  RPKDPEGVPPLLVSQQ | 1 | 0.929698 | 0.4503 | 0.5711 |
| ALK tyrosine kinase receptor | https://www.iedb.org/epitope/2257725 | PEGVPPLLVSQQAKREEERSP  AAPPPLPTTSSGKAA | 1 | 0.962538 | 0.4698 | 0.7611 |
| ALK tyrosine kinase receptor | https://www.iedb.org/epitope/2257726 | PPLPTTSSGKAAKKPTAAEISV  RVPRGPAVEGGHVN | 1 | 0.47973 | 0.4681 | 0.7855 |
| https://www.iedb.org/epitope/1311443 | SMAD5 antisense gene protein 1 | PPPPPKAPAGQETLSLQSR | 1 | 0.616362 | 0.4486 | 0.4008 |
| Mucin-2 | https://www.iedb.org/epitope/2232836 | PPTTTPSPPPTSTTTLPPT | 1 | 0.63489 | 0.3673 | 0.405 |
| ALK tyrosine kinase receptor | https://www.iedb.org/epitope/2257727 | PRGPAVEGGHVNMAFSQSN  PPSELHKVHGSRNKPTS | 1 | 0.558497 | 0.4914 | 0.7479 |
| Folate receptor alpha | https://www.iedb.org/epitope/2238425 | PWAAWPFLLSLALMLLWL | 1 | 0.89605 | 0.4566 | 0.33 |
| Minor capsid protein L2 | https://www.iedb.org/epitope/110385 | QLYKTCKQAGTCPPDIIPKV | 1 | 0.923137 | 0.4524 | 0.5383 |
| Melanoma-associated antigen 3 | https://www.iedb.org/epitope/2275808 | QRSQHCKPEEGLEARGEAL | 1 | 0.553437 | 0.4853 | 0.3888 |
| Transcription factor SOX-2 | https://www.iedb.org/epitope/1346775 | RALHMKEHPDYKYRPRRKT  KTLMKKDK | 1 | 0.972722 | 0.4337 | 0.7239 |
| Endogenous retrovirus group K member 18 Env polyprotein | https://www.iedb.org/epitope/1863649 | RPKGKTCPKEIPKGSKNT | 1 | 0.914791 | 0.4504 | 0.6753 |
| Folate receptor alpha | https://www.iedb.org/epitope/2238445 | RTELLNVCMNAKHHKEK | 1 | 0.972846 | 0.4724 | 0.249 |
| ALK tyrosine kinase receptor | https://www.iedb.org/epitope/2257730 | SCTVPPNVATGRLPGASLLL  EPSSLTANMKEVPLFR | 1 | 0.369105 | 0.4993 | 0.6071 |
| ALK tyrosine kinase receptor | https://www.iedb.org/epitope/2257731 | SDLKEVPRKNITLIRGLGHG  AFGEVYEGQVSGMPND | 1 | 0.220866 | 0.4799 | 0.8356 |
| Putative uncharacterized protein encoded by LINC00615 | https://www.iedb.org/epitope/1311493 | SFFFFKQPSPKTQDHLRQQRASAR | 1 | 0.601364 | 0.498 | 0.6358 |
| Transcription factor SOX-2 | https://www.iedb.org/epitope/1346783 | SRGQRRKMAQENPKMHNSE  ISKRLGAEWKLLSETEK | 1 | 0.640853 | 0.5298 | 0.8054 |
| Myristoylated alanine-rich C-kinase substrate | https://www.iedb.org/epitope/1311512 | TPKKKEALFLQEVFQAERLLLQEEQEGGWRRR | 1 | 0.904065 | 0.4956 | 0.6731 |
| Myristoylated alanine-rich C-kinase substrate | https://www.iedb.org/epitope/1311513 | TPKKKRSAFPSRSLSS | 1 | 0.523728 | 0.4908 | 0.5811 |
| Receptor tyrosine-protein kinase erbB-2 | https://www.iedb.org/epitope/2237943 | VARCPSGVKPDLSYMPIWKFPDEEGACQPL | 1 | 0.508183 | 0.4396 | 0.6836 |
| Activin receptor type-2A | https://www.iedb.org/epitope/1311517 | VHKKEACFKRLLAETCWNGNAL | 1 | 0.666383 | 0.4919 | 0.4185 |
| Endogenous retrovirus group K member 18 Env polyprotein | https://www.iedb.org/epitope/1711669 | VWVPGPTDDRCPAKPEEEG | 1 | 0.954102 | 0.4317 | 0.2154 |
| ALK tyrosine kinase receptor | http://iedb.org/epitope/2257738 | VYEGQVSGMPNDPSPLQ  VAVKTLPEVCSEQDELDFL | 1 | 0.885815 | 0.4958 | 0.8136 |
| Tyrosine-protein kinase Lck | http://www.iedb.org/epitope/2266761 | ASPLQDNLVIALHSY | 0 | 0.047313 | 0.4409 | 0.3807 |
| Tyrosine-protein kinase Lck | http://www.iedb.org/epitope/2266772 | DFGLARLIEDNEYTA | 0 | 0.202336 | 0.4395 | 0.6654 |
| lymphocyte-specific protein tyrosine kinase | http://www.iedb.org/epitope/2266773 | DGDLGFEKGEPLRIL | 0 | 0.344582 | 0.4528 | 0.5061 |
| nuclease-sensitive element-binding protein 1 [Homo sapiens] | http://www.iedb.org/epitope/754695 | DGETVEFDVVEGEKG | 0 | 0.022651 | 0.4597 | 0.5384 |
| Tyrosine-protein kinase Lck | http://www.iedb.org/epitope/2266777 | DNLVIALHSYEPSHD | 0 | 0.28952 | 0.4442 | 0.482 |
| nuclease-sensitive element-binding protein 1 [Homo sapiens] | http://www.iedb.org/epitope/754790 | DRNHYRRYPRRRGPP | 0 | 0.593243 | 0.3681 | 0.7247 |
| nuclease-sensitive element-binding protein 1 [Homo sapiens] | http://www.iedb.org/epitope/754845 | EAANVTGPGGVPVQG | 0 | 0.002823 | 0.4201 | 0.6181 |
| nuclease-sensitive element-binding protein 1 [Homo sapiens] | http://www.iedb.org/epitope/754877 | EDGNEEDKENQGDET | 0 | 0.001681 | 0.4621 | 0.7413 |
| nuclease-sensitive element-binding protein 1 [Homo sapiens] | http://www.iedb.org/epitope/754892 | EEDKENQGDETQGQQ | 0 | 0.000518 | 0.4825 | 0.6261 |
| nuclease-sensitive element-binding protein 1 [Homo sapiens] | http://www.iedb.org/epitope/754948 | EKGAEAANVTGPGGV | 0 | 0.003024 | 0.459 | 0.6166 |
| nuclease-sensitive element-binding protein 1 [Homo sapiens] | http://www.iedb.org/epitope/754955 | EKNEGSESAPEGQAQ | 0 | 0.004645 | 0.4863 | 0.4668 |
| nuclease-sensitive element-binding protein 1 [Homo sapiens] | http://www.iedb.org/epitope/754990 | ENPKPQDGKETKAAD | 0 | 0.031971 | 0.4286 | 0.5328 |
| nuclease-sensitive element-binding protein 1 [Homo sapiens] | http://www.iedb.org/epitope/754992 | ENQGDETQGQQPPQR | 0 | 0.058405 | 0.4621 | 0.595 |
| Tyrosine-protein kinase Lck | http://www.iedb.org/epitope/2266786 | FDYLRSVLEDFFTAT | 0 | 0.207217 | 0.4519 | 0.2025 |
| Tyrosine-protein kinase Lck | http://www.iedb.org/epitope/2266787 | FIPFNFVAKANSLEP | 0 | 0.219834 | 0.4594 | 0.4534 |
| nuclease-sensitive element-binding protein 1 [Homo sapiens] | http://www.iedb.org/epitope/755136 | FNYRRRRPENPKPQD | 0 | 0.145213 | 0.3946 | 0.778 |
| nuclease-sensitive element-binding protein 1 [Homo sapiens] | http://www.iedb.org/epitope/755155 | FRRGPPRQRQPREDG | 0 | 0.300872 | 0.4238 | 0.4528 |
| nuclease-sensitive element-binding protein 1 [Homo sapiens] | http://www.iedb.org/epitope/755179 | GAGSGGPGGLTSAAP | 0 | 0.027718 | 0.4055 | 0.7043 |
| Tyrosine-protein kinase Lck | http://www.iedb.org/epitope/2266791 | GEVVKHYKIRNLDNG | 0 | 0.200951 | 0.4617 | 0.6873 |
| nuclease-sensitive element-binding protein 1 [Homo sapiens] | http://www.iedb.org/epitope/755234 | GFINRNDTKEDVFVH | 0 | 0.358108 | 0.4653 | 0.5072 |
| nuclease-sensitive element-binding protein 1 [Homo sapiens] | http://www.iedb.org/epitope/755247 | GGPGGLTSAAPAGGD | 0 | 0.007754 | 0.421 | 0.8133 |
| nuclease-sensitive element-binding protein 1 [Homo sapiens] | http://www.iedb.org/epitope/755255 | GGVPVQGSKYAADRN | 0 | 0.452295 | 0.4431 | 0.445 |
| nuclease-sensitive element-binding protein 1 [Homo sapiens] | http://www.iedb.org/epitope/755302 | GLTSAAPAGGDKKVI | 0 | 0.01964 | 0.4215 | 0.779 |
| nuclease-sensitive element-binding protein 1 [Homo sapiens] | http://www.iedb.org/epitope/755337 | GPPRNYQQNYQNSES | 0 | 0.00601 | 0.4571 | 0.451 |
| nuclease-sensitive element-binding protein 1 [Homo sapiens] | http://www.iedb.org/epitope/755389 | GSESAPEGQAQQRRP | 0 | 0.014439 | 0.4585 | 0.5911 |
| Tyrosine-protein kinase Lck | http://www.iedb.org/epitope/2266796 | GSFLIRESESTAGSF | 0 | 0.033763 | 0.4937 | 0.3252 |
| nuclease-sensitive element-binding protein 1 [Homo sapiens] | http://www.iedb.org/epitope/755428 | GTVKWFNVRNGYGFI | 0 | 0.418672 | 0.4653 | 0.483 |
| Tyrosine-protein kinase Lck | http://www.iedb.org/epitope/2266804 | INKLLDMAAQIAEGM | 0 | 0.070188 | 0.4695 | 0.5206 |
| Tyrosine-protein kinase Lck | http://www.iedb.org/epitope/2266806 | IYIITEYMENGSLVD | 0 | 0.034968 | 0.4806 | 0.3298 |
| nuclease-sensitive element-binding protein 1 [Homo sapiens] | http://www.iedb.org/epitope/755736 | KEDVFVHQTAIKKNN | 0 | 0.640794 | 0.4867 | 0.563 |
| nuclease-sensitive element-binding protein 1 [Homo sapiens] | http://www.iedb.org/epitope/755845 | KNNPRKYLRSVGDGE | 0 | 0.322311 | 0.4496 | 0.4045 |
